# DESIGN AND GENERATION OF A RECOMBINANT GAMMAHERPESVIRUS ENCODING shRNA FROM NATIVE VIRAL tRNA PROMOTER

**DOI:** 10.1101/2023.02.02.523555

**Authors:** Mehmet Kara, Scott Tibbetts

## Abstract

Gammaherpesviruses are associated with multiple types of tumour development and understanding the pathogenesis of these viruses has been the subject of many different studies. Throughout the lytic and latent life cycle, these viruses utilize numerous virally encoded miRNAs to regulate the key mechanisms of the infected cell in their favour. Therefore it is important to understand the miRNA and their mRNA target interactions for developing better therapeutics. In this study, the strategy and design of a short hairpin encoding RNA element targeting Blimp1 transcript in the context of viral infection is evaluated. This proof of principle experiment provides a mean to study important miRNA mRNA interactions *in vivo*. The very short promoter size of the Murine gammaherpesvirus 68 (MHV68) vtRNA4 and its ability to generate two shRNAs from a ~180 nucleotide sequence is useful if there is a size limit for the shRNA construct.

## Introduction

Gammaherpesvirus infections are highly prevalent in human populations and persistent for the life of those hosts with a functional immune system (Arvin *et al*., 2007). The persistent infection is called latency, and is characterized by expression of a limited set of viral genes and noncoding RNAs, including miRNAs (Speck & Ganem, 2010). Chronic gammaherpesvirus infections are associated with the development of lymphoproliferative diseases and certain types of lymphomas such as diffuse large B cell lymphoma in individuals with immune deficiencies, including those with primary immunodeficiencies, HIV infection, and organ transplant patients under immune suppression (Barton *et al*., 2011). Thus, understanding the establishment and maintenance of latency programs is important for developing preventative strategies and treatments for gammaherpesvirus pathogenesis as well designing gene delivery vectors and vaccines.

One of the key regulators of herpesvirus latency is virally encoded miRNAs (Cullen, 2011). miRNAs are short, 20-22 nucleotide long RNA molecules that play a key role in post transcriptional gene regulation (Bartel, 2009). Up to date, many viral miRNAs have been identified within different viral families including gammaherpesvirus subfamily (Carnero *et al*., 2011; Jurak *et al*., 2011; Pfeffer *et al*., 2004; Samols *et al*., 2005). Due to the narrow host range of human gammaherpesviruses such as Kaposi’s Sarcoma Associated Herpesvirus (KSHV) and Epstein Barr Virus (EBV), the murine gammaherpesvirus 68 (MHV68) offers a small animal model infection system to study viral pathogenesis *in vivo* (Simas & Efstathiou, 1998). In recent years, several studies identified the “targetome” of viral miRNAs by utilizing Ago protein immunoprecipitation-linked sequencing based methods (Bullard *et al*., 2019; Chi *et al*., 2009; Helwak *et al*., 2013). Similar to the human gammaherpesviruses, MHV68 encodes multiple miRNAs (Reese *et al*., 2010; Zhu *et al*., 2010). Interestingly though, MHV68 miRNAs are transcribed from viral tRNA promoters as approximately 200 nucleotide long tRNA-miRNA linked elements called TMERs and processed to become mature miRNAs through tRNase Z and Dicer (Bogerd *et al*., 2010).

The biological importance of a given viral miRNA and its target mRNA or lncRNA is mostly understudied and requires new and applicable methods. In order to study viral miRNA-mRNA interactions in an animal model it is hypothesized target mRNA specific short hairpin RNAs (shRNA) can be used to understand the function of a specific miRNA and mRNA interaction *in vivo*. Since miRNAs can bind and repress more than one target gene, shRNA strategy will specifically pinpoint the function of a given miRNA-mRNA interaction.

Generally, shRNAs are designed to produce a 21-22 nucleotide long RNA molecule that is perfectly complementary to the target mRNA while miRNAs bind to their targets with imperfect complementarity. shRNAs are most commonly transcribed by a U6 promoter, which is an RNA III dependent promoter, from a plasmid vector (Moore *et al*., 2010). In this study we have tested whether a virally encoded tRNA, that also consists of an RNA polymerase III promoter, can drive the expression of an shRNA and whether the designed shRNAs are functional. Additionally, we have generated a mutant virus that expresses the designed shRNA from a neutral locus within the viral genome. The shRNAs used in this study were designed for mouse *Prdm1* gene product. *Prdm1* gene codes for Blimp1 protein which is a transcription factor that drives the transition of B cells from a germinal center (GC) phenotype to a plasma cell phenotype by repressing the expression of GC-specific genes (Calame, 2006; Kallies & Nutt, 2007). Since MHV68 naturally establishes latency in B cells, this mutant virus becomes a valuable tool to investigate miRNA-mRNA interactions as a proof of concept.

## Material and Methods

### Desing and synthesis of the shRNAs

MHV68 viral tRNA4 naturally contains two stem loops for miRNAs and we designed two shRNA hairpins. The first one of the Blimp1 shRNAs was designed using the Invivogen siWizard algorithm (https://www.invivogen.com/sirnawizard/construct.php) (*SiRNA Wizard - Design Hairpin Insert - InvivoGen*, n.d.) by using standard criteria offered by the website. The other shRNA sequence was designed by using sequence information of an siRNA that is used and successfully knocked down Blimp1 in a previously published paper (Zhou *et al*., 2013). siWizard is also used to design the scrambled version of these shRNAs. Then these sequences are inserted at nucleotide position 74 of vtRNA4 based on where the naturally occurring miRNA starts and the in silico folding analysis of vtRNA4 and its miRNAs. The shRNA sequences were separated by a spacer sequence (TTACGT) and inserted in place of viral miRNAs encoded by TMER4. The *in silico* folding analysis of the end construct was performed using the mFOLD algorithm (http://www.unafold.org/RNA_form.php) (*RNA Folding Form for 88.230.102.221*, n.d.). The Blimp1 shRNAs and control shRNAs scrambled constructs were synthetically produced by Genewiz. Primers and shRNA/scrRNA sequences are listed in Table 1.

**Table 1.**
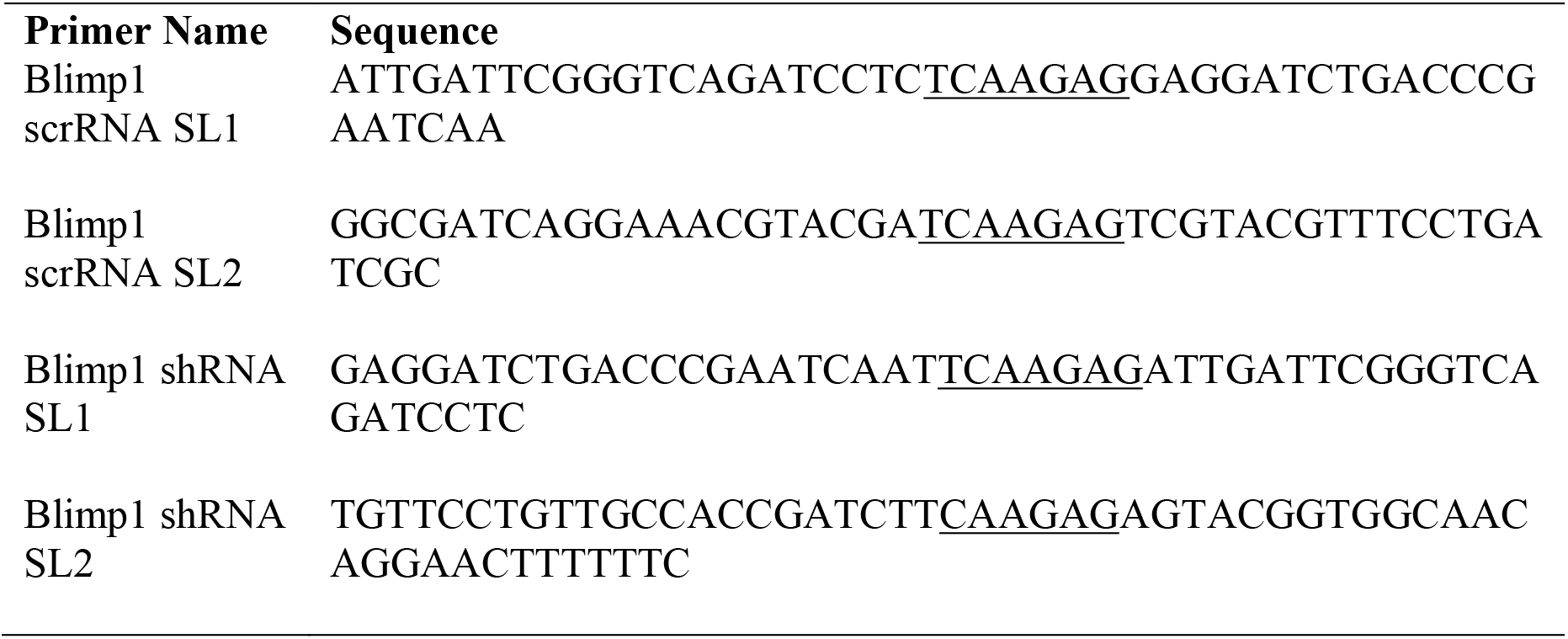
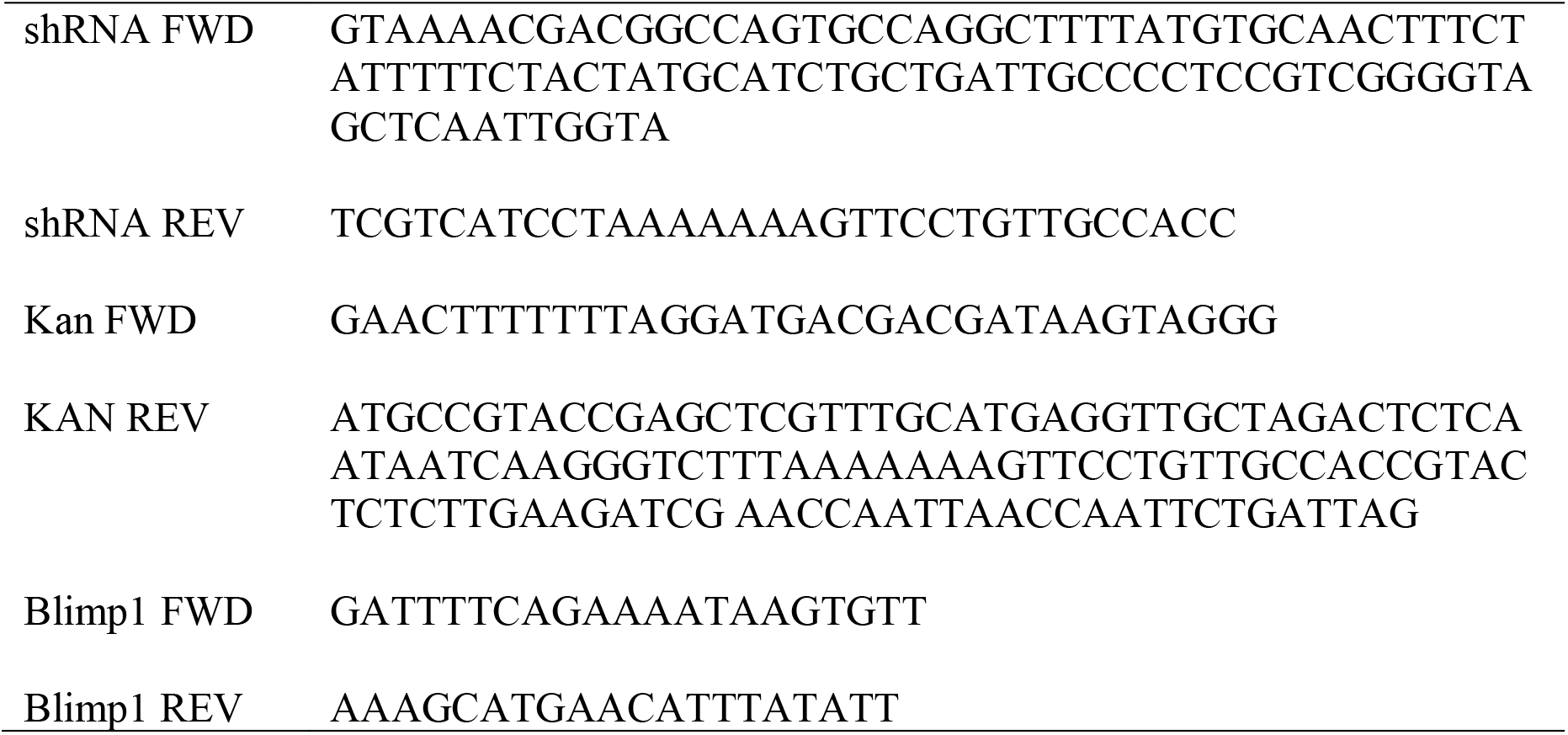
List of primers and shRNA sequences. The underlined sequences are the loop sequences of the stem loops in shRNA designs.

### Generation of shRNA-Encoding Plasmids and Viruses

The ~200 base pair long sequences were PCR-amplified and cloned with a Kanamycin resistance PCR fragment into pUC19 vector by using GeneArt Seamless Cloning Assembly kit (Life Technologies) to generate the targeting construct. The recombinant viruses MHV68.Blimp1-sh and MHV68.Blimp1-scr were generated in two stages using two-step Red-mediated recombination onto the wild-type MHV68.ORF73βla BAC backbone (Nealy *et al*., 2010). The shRNA-Kanamycin fragments were then amplified from the plasmids, with PCR amplicons containing homologous regions of upstream and downstream of the insertion site, which are required for intramolecular recombination. The resulting amplicon was gel-purified, then electroporated into GS1783 *E. coli* cells containing the MHV68.ORF73βla BAC and Red recombinase machinery (Tischer *et al*., 2006). Transformed cells were incubated at 30°C overnight on Kan+ agar plates. DNA from the resulting colonies was isolated and digested with XhoI, then screened for the insertion genomic integrity by gel electrophoresis (GE). In selected positive clones, I-Sce was induced by incubation for 2 h in 1% arabinose to induce Red recombinase expression for the second recombination. Resulting colonies were again screened by PCR and GE, then exact sequence was confirmed by Sanger sequencing. BAC DNA from the positive clones were isolated with a Qiagen large-construct plasmid purification kit, then transfected into NIH 3T12 cells using a TransIT-3T3 transfection kit (MirusBio). A GFP cassette in the BAC DNA was used to visualize the transfection and propagation of the virus. To excise the BAC cassette, the resulting virus from the initial transfection was used to infect NIH 3T12 cells expressing Cre recombinase at an Multiplicity of Infection (MOI) of 0.05, and passaged twice. Titers of final viral BAC-minus virus stocks were determined by plaque assay on NIH 3T12 cells.

### Luciferase Assays

The pGL3-promoter vector from Promega was used for firefly luciferase assays. pGL3-promoter vector was digested with Xba I. Putative mRNA target regions were PCR-amplified to contain the Xba I site and homology to the pGL3-promoter, and then were incubated with linearized pGL3-promoter vector and the Gibson Assembly Kit reagents (NEB). Reactions were then transformed into competent Top10 *E. coli*. The transformants were then Sanger sequenced to validate insertion of the target sequence into the 3’ UTR of the pGL3-promoter vector. For transfections, 10,000 NIH 3T12 fibroblasts were seeded onto 96-well plates the day before transfection. For shRNA and TMER plasmids, 1ng of pRL-SV40 renilla luciferase vector, 100ng of the pGL3-promoter containing the target sequence in the 3’ UTR, and 100ng of the TMER or shRNA vector was transfected with Lipofectamine 2000 (Thermofisher). After 48 hours the efficiency was checked visually by GFP control plasmid transfection, then supernatant fluid was removed from wells, and 50μL of fresh DMEM plus 50 μL of Dual Glo Luciferase Reagent (Promega) was added back. Samples were incubated for 10 minutes, then transferred to 96-well NUNC plates, and firefly luciferase was measured using a plate luminometer. Next, 50 μL of Dual Glo Stop Reagent was added, and renilla luciferase activity was measured. Renilla luciferase activity was used to control for transfection efficiency. For the infection and transfection luciferase assays, virus transfections are done in 12 well plates. 10^5^ NIH 3T12 cells were plated into each well of a 12 well plate the day before infection/transfection. For infection first scheme, cells are washed with PBS and infected at multiplicity of infection (MOI) of 5 with 200 μL of media without FBS and incubated at 37C for 1 hour with gentle mixing every 15 minutes. Then the infection media is removed washed with PBS once and incubated with 400 μL of plasmid lipofectamine mixture for 4 hours at 37C. at the end of the incubation period 1mL media is added to each well. If the cells are transfected first, transfection was conducted first as and then cells were infected mentioned above. 48 hours post infection cells are assayed for luciferase activity as described in above. Data were analyzed by either Excel or Graphpad Prism software.

## Results

The majority of the MHV68 encoded **t**RNA-**m**iRNA **E**ncoding **R**NAs (TMERs) contain two stem loops which each are capable of generating two mature miRNAs. TMER4 expresses mghv-miR-M1-5-5p and mghv-miR-M1-6-3p from stem loops 1 and 2, respectively. mghv-miR-M1-5-5p is expressed in both lytically-infected fibroblasts and latent cell lines, and in latently-infected mouse splenocytes *in vivo*. mghv-miR-M1-6-3p is expressed at very low levels in tissue culture, but readily detected in latently infected mouse splenocytes in vivo (Feldman *et al*., 2014) For this reason, the TMER4 promoter element within vtRNA4 was chosen to drive expression of the the *Blimp1* targeting-shRNAs. One of the shRNAs was designed by using an online algorithm SiWizard (Invivogen) and the other shRNA sequence was previously shown to inhibit Blimp1 expression *in vivo* ((Zhou *et al*., 2013). In order to ensure the proper folding and structure of the construct, the sequences are folded *in silico* by mFOLD (Figure 1A). The natural miRNA secondary RNA structure shows bulges in the hairpin region whereas shRNA hairpins are perfectly complementary as expected and there was no alternation in the predicted tRNA folding with the exchange of sequences (Figure 1B). The Blimp1 targeting shRNA is tested by luciferase assay and compared to the scrambled RNA for the ability to bind and inhibit the 3’ UTR of Blimp1 transcript cloned into the luciferase vector pGL3. The combination of two shRNAs showed approximately 65% of reduction in luciferase assay (Figure 1C).

**Figure 1.**
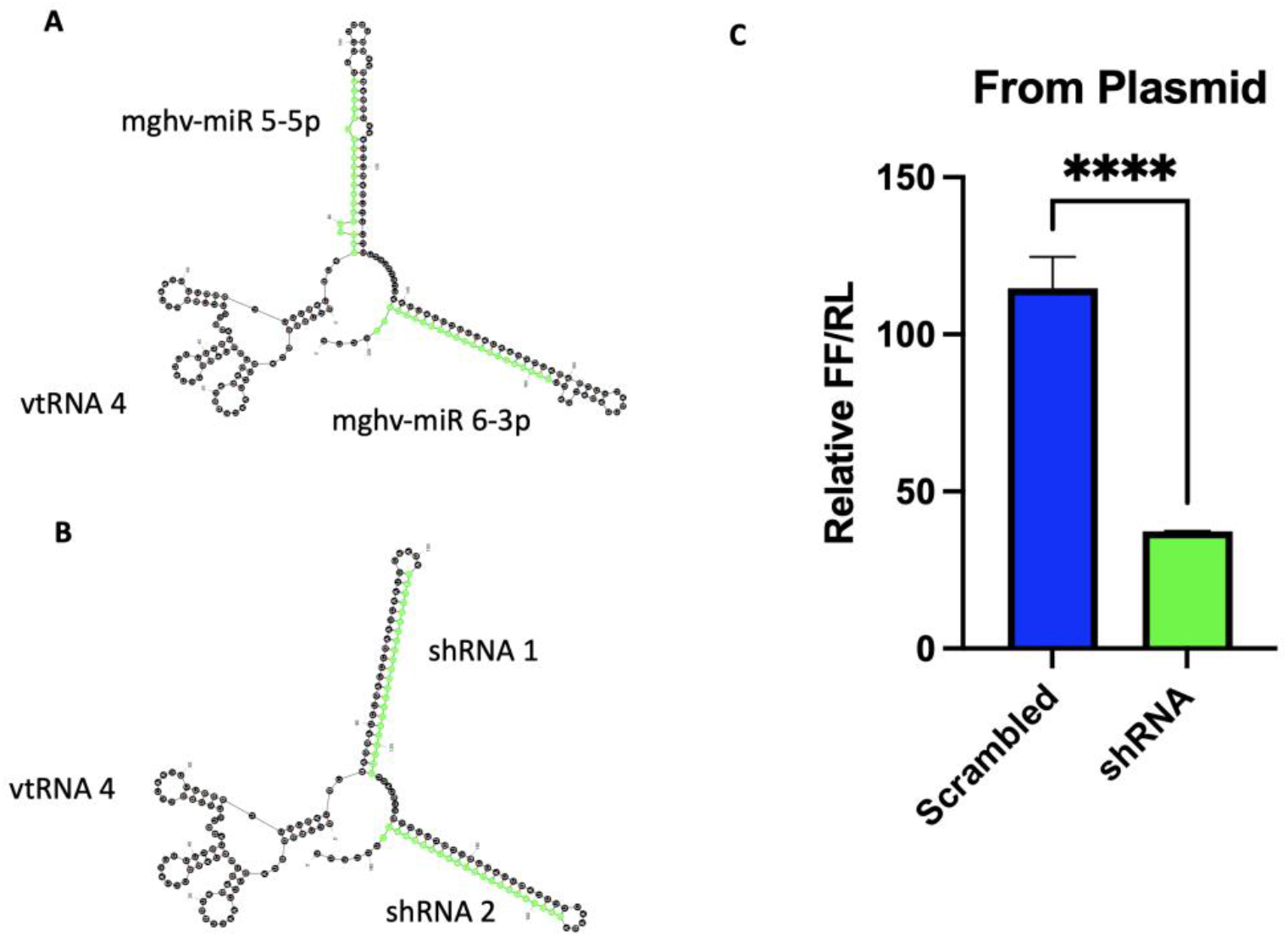
Design and testing of Blimp1 shRNA and scrambled control. A) MHV68 vtRNA4 and corresponding miRNA stemloop sequences in mfold secondary structure prediction. B) vtRNA and two Blimp1 targeting shRNA construct RNA folding prediction in mfold. C) Luciferase assay. shRNA target region containing firefly luciferase vector pGL3, renilla luciferase vector and shRNA vector or its scrambled control is transfected to NIH3T12 cells and assayed for luciferase. Relative values of firefly over renilla luciferase activity are shown. ****P value <0.001.

Since the shRNA constructs worked efficiently, we then proceed to generation of the shRNA encoding virus. Briefly, the targeting construct containing a kanamycin resistance cassette and the shRNA or scrambled version was prepared and inserted between TMER1 and TMER2 by Red mediated mutagenesis protocol. The resulting viral clones are screened and as shown in the gel electrophoresis for the viral BAC integrity (Figure 2). The intact clones are selected and viral stocks are prepared after transfecting BAC DNA into 3T12 cells and passaging the virus through Cre recombinase expressing 3T12 cells to remove the BAC cassette from the virus genome.

**Figure 2.**
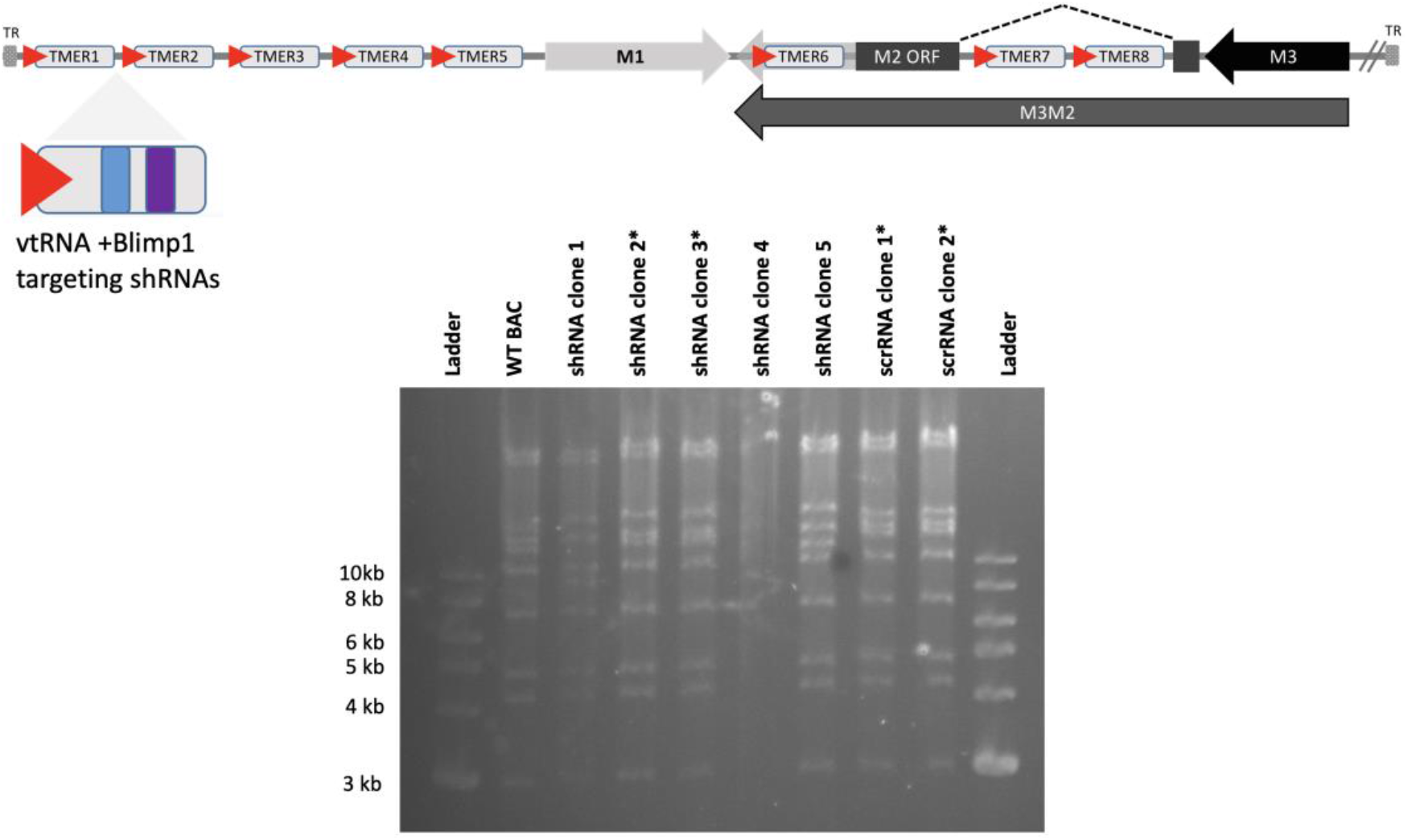
Schematic representation of left end of the viral genome and screening of recombinant clones. Upper part of figure displays the relative insertion site for the shRNA or scrRNA constructs. vtRNA is viral tRNA. Red arrow is the promoter for tRNA and blue and purple boxes are shRNA stem loops 1 and 2 respectively. After Red recombination the mutant clones are screened for integrity of the virus and compared to wild type (WT). The ones with * are the correct clones used to generate viral stocks.

After the virus stocks were prepared and tittered, we have tested the shRNA construct in the context of viral infection. It is expected that the shRNAs are expressed from the virus however, all the other viral components produced from the virus has to be evaluated. It is conceivable that cells are stressed because the viral infection and transfection are simultaneously happening. Therefore we tested two different options for infection first and then transfection as well as the vice versa. Even though to a lesser degree when compared to the plasmid produced shRNA, the virally encoded shRNA still showed a 50% reduction in the luciferase activity (Figure 3). One possible explanation for the lesser degree reduction in the viral infection versus the plasmid transfection (Figure 1C and Figure 3) is that during viral infection many more miRNAs are present from the virus in the cell, therefore less RISC processing machinery is available for the shRNAs.

**Figure 3.**
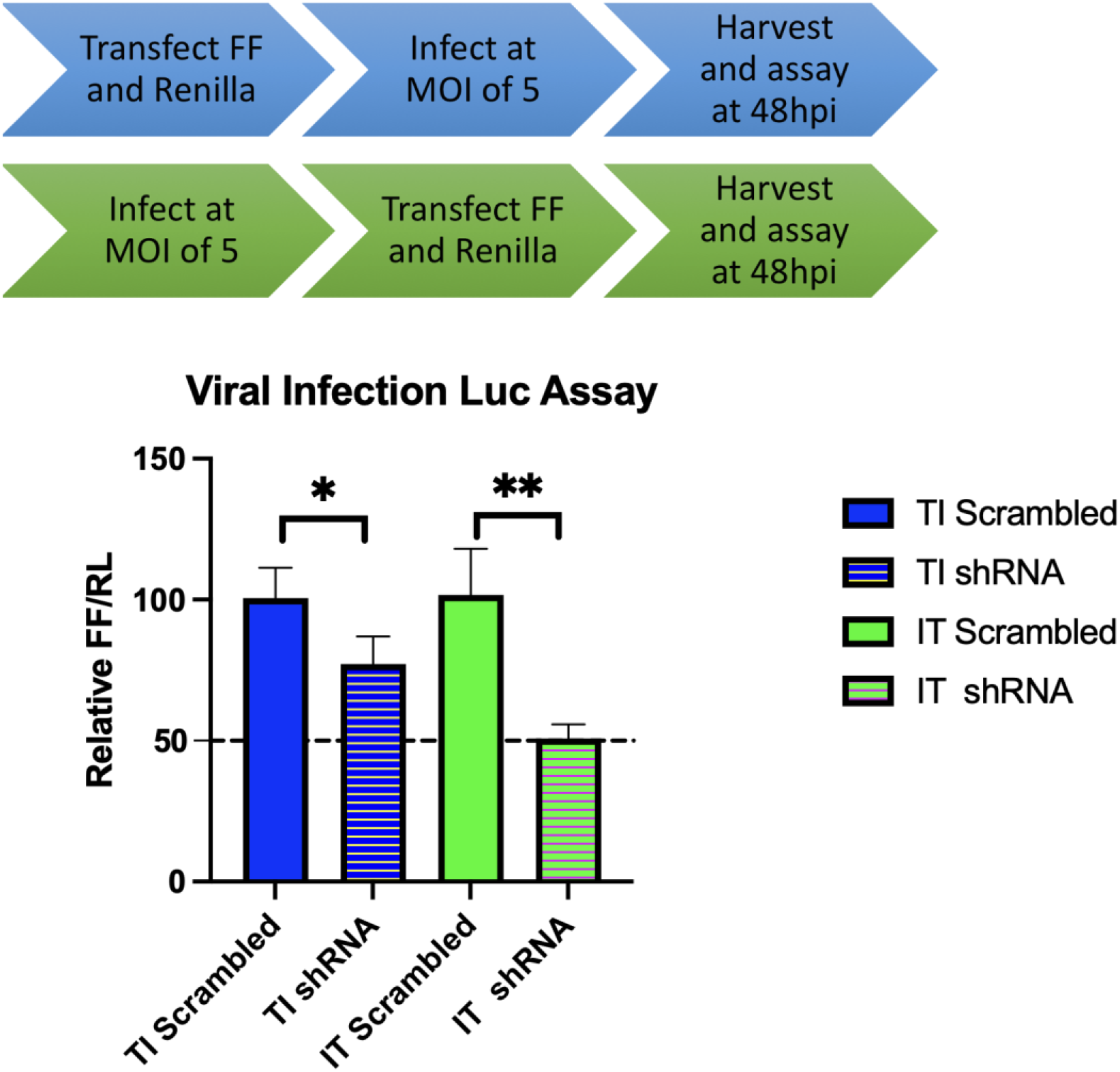
Transfection and infection scheme and luciferase assay. Infection first transfection second or vice versa approach is used to test shRNA constructs. FF is firefly luciferase vector containing the shRNA target sequence. MOI indicates the multiplicity of infection. 5 plaque forming unit (pfu) virus used per cell in the assay and luciferase readings are recorded at 48 hours post infections (hpi). Y axis grids indicated the 100% and 50%. Data is displayed as relative values of firefly luciferase over renilla luciferase activity. TI is transfection infection, IT is infection transfection. P value * 0.0186, ** 0,0019

## Discussion and Conclusion

Defining the biological role of viral miRNA targeting of a transcript *in vivo* is one of the key aspects to study miRNA function in a relevant pathogenesis model. One means to examine the role of regulation of a specific target transcript (as opposed to other targets of the same miRNA) is to knock down that transcript more precisely using shRNAs, ideally expressed from the virus in place of the miRNA in question. In order to test this possibility, we have designed a proof-of-concept experiment to show whether shRNAs generated from the murine gammaherpesvirus 68 encoded TMER constructs are functional. To test this, we have designed shRNAs targeting the *Prdm1* gene that produces Blimp1 protein. Blimp1 is a key regulator of the germinal center B cell transition to memory B cells. (Calame, 2006; Kallies and Nutt, 2007). Therefore in theory, *Prdm1*-targeting shRNAs encoded by the virus would result in down regulation of the Blimp1 protein, and preventing differentiation of the infected cell into an antibody-secreting plasma cell.

This construct containing the TMER4 promoter and two shRNA stem loops was then tested for its ability to knock down *Prdm1* expression in luciferase assays. The TMER4 promoter shRNA-encoding element was able to knock down luciferase construct carrying Blimp1 sequence, showing that (a) functional shRNAs were produced under control of the vtRNA promoter, and (b) the shRNAs could successfully knock down the target transcript. By using two-step red recombination, then generated an shRNA-encoding virus in which the TMER 4 promoter shRNA cassette was inserted in a ‘neutral’ region between TMER1 and TMER2. This virus and its control were also tested in luciferase assays.The sequential transfection and infection versus infection and transfection was compared. In both conditions, the shRNA-encoding virus displayed knock down of the target; however, infecting cells before transfection worked with better efficiency. The possible explanation for the difference in these two schematic is that the cells more readily take up plasmids after infection and the virus probably starts making shRNAs from the very early time points. However, if cells are transfected first, it is more likely that the infection ability of the virus is reduced because the cells are already stressed because of the transfection reagents and DNA and therefore less shRNAs are produced by the virus.

One of the contributions of this study is the design of a viral tRNA promoter for shRNAs as an alternative to commonly used U6 and H1 RNA polymerase III promoters which are around 250 to 350 nucleotides only for the promoter sequence. Usage of the tRNA promoters have been tested before for the generation of shRNAs. tRNA^Lys3^ promoter was used to deliver HIV tat/rev shRNA successfully (Scherer *et al*., 2007). Here we have utilized a similar strategy for one of the naturally occurring MHV68 vtRNAs, vtRNA4 which is only 74 nucleotides long. If the shRNA sequence is added before CCA addition site of this viral tRNA, a very minimal RNA pol III promoter can be used to express shRNAs. In the context of MHV86 vtRNAs, most of them can be utilized to deliver at least two hairpins if a stronger knockdown is desired. From a 180 nucleotide construct (promoter + two shRNA stem loops) a desired gene can be knock down in the context of virus infection in a given specific cell (for example B cells for gammaherpesviruses) *in vivo*.

## Acknowledgements

I would like to thank Tibbetts’ Lab members for their valuable comments. This work was conducted during author’s PhD studies.

## Ethical Statement

This study did not require an ethical committee approval.

